# Admixture as a source for HLA variation in Neolithic European farming communities

**DOI:** 10.1101/2023.08.23.554285

**Authors:** Nicolas Antonio da Silva, Onur Özer, Magdalena Haller, Yan-Rong Chen, Daniel Kolbe, Sabine Schade-Lindig, Joachim Wahl, Carola Berszin, Michael Francken, Irina Görner, Kerstin Schierhold, Joachim Pechtl, Gisela Grupe, Christoph Rinne, Johannes Müller, Tobias L. Lenz, Almut Nebel, Ben Krause-Kyora

## Abstract

The northern European Neolithic is characterized by two major demographic events: immigration of early farmers (EF) from Anatolia (5500 BCE) and their admixture (from ∼4200 BCE) with western hunter-gatherers (WHG) forming late farmers (LF). The influence of this admixture event on variation in the immune-relevant human leukocyte antigen (HLA) region is understudied. Here, we conducted population and immunogenetic analyses on 83 individuals from six EF and LF sites located in present-day Germany. We observed significant shifts in HLA allele frequencies from EF to LF. The HLA diversity increased from EF to LF, likely due to admixture with WHG. However, it was considerably lower than in modern populations. Both EF and LF exhibited a relatively narrow HLA allele spectrum compared to today. This coincides with sparse traces of pathogen DNA, potentially indicating a lower pathogen pressure at the time. We additionally noted that LF resulted from sex-biased admixture from male WHG.

**TEASER:** More restricted HLA allele spectrum and lower diversity in Neolithic farmers than in modern populations

## INTRODUCTION

Since the Palaeolithic, central Europe had been populated by western hunter-gatherers (WHG). Around 5500 BCE, the first farmers arrived who originated from Anatolia, bringing with them agriculture as subsistence and the Neolithic lifestyle (1). Archaeologically, these early European farmers are associated with the Linear Pottery societies (Linearbandkeramik, LBK, ∼5500–4900 BCE). LBK and subsequent societies remained largely unadmixed with WHG, as reflected in their high genetic similarity to the Anatolian source populations (2, 3). The rate of admixture gradually increased from the Younger and Late Neolithic (4200–2800 BCE) onwards, so that the gene pool of the resulting late farmers contained a substantial WHG ancestry component (2–5). These demographic and genomic changes coincided with cultural transformations that led to the dissolution of LBK/post-LBK societies and ultimately to the emergence of many small and regionally diverse societies, such as the one affiliated with the Wartberg context (WBC, ∼3500– 2800 BCE) (5–7). So far, only one WBC burial community (i.e., Niedertiefenbach, 3300–3200 BCE) has been comprehensively studied by ancient genomics (5). This group had a surprisingly high WHG ancestry (34-58%) and a distinct human leukocyte antigen (HLA) immune gene profile that was mainly focused on the detection of viral infections (5). However, whether these genomic characteristics of the Niedertiefenbach population were typical of the WBC in general remains to be clarified. Another question is to what extent the HLA repertoire of the WBC-associated farmers differed from that of earlier groups, for instance, the LBK and post-LBK communities.

HLA molecules play a key role in adaptive immunity and exhibit exceptional levels of polymorphism, presumably driven by pathogen-mediated selection (8–10). Improved ancient DNA (aDNA) technology has recently yielded the first prehistorical studies on HLA alleles obtained after sequencing the HLA region, providing initial glimpses into the co-evolutionary history of humans and their pathogens (5, 11).

Here, we performed population genetic analyses using newly generated genome-wide data of 83 individuals from six archaeological sites covering the 2300-year transition from the Early (LBK) to the Late Neolithic (WBC) (Figs. 1A-1B, Table 1). Moreover, we sequenced 80 individuals for HLA variation (Table 1), thus significantly expanding the publicly available data (5, 11, 12). This larger sample size provided more reliable HLA allele frequencies and allowed us to perform more robust and informative comparisons between Neolithic and modern populations.

**Fig. 1.**
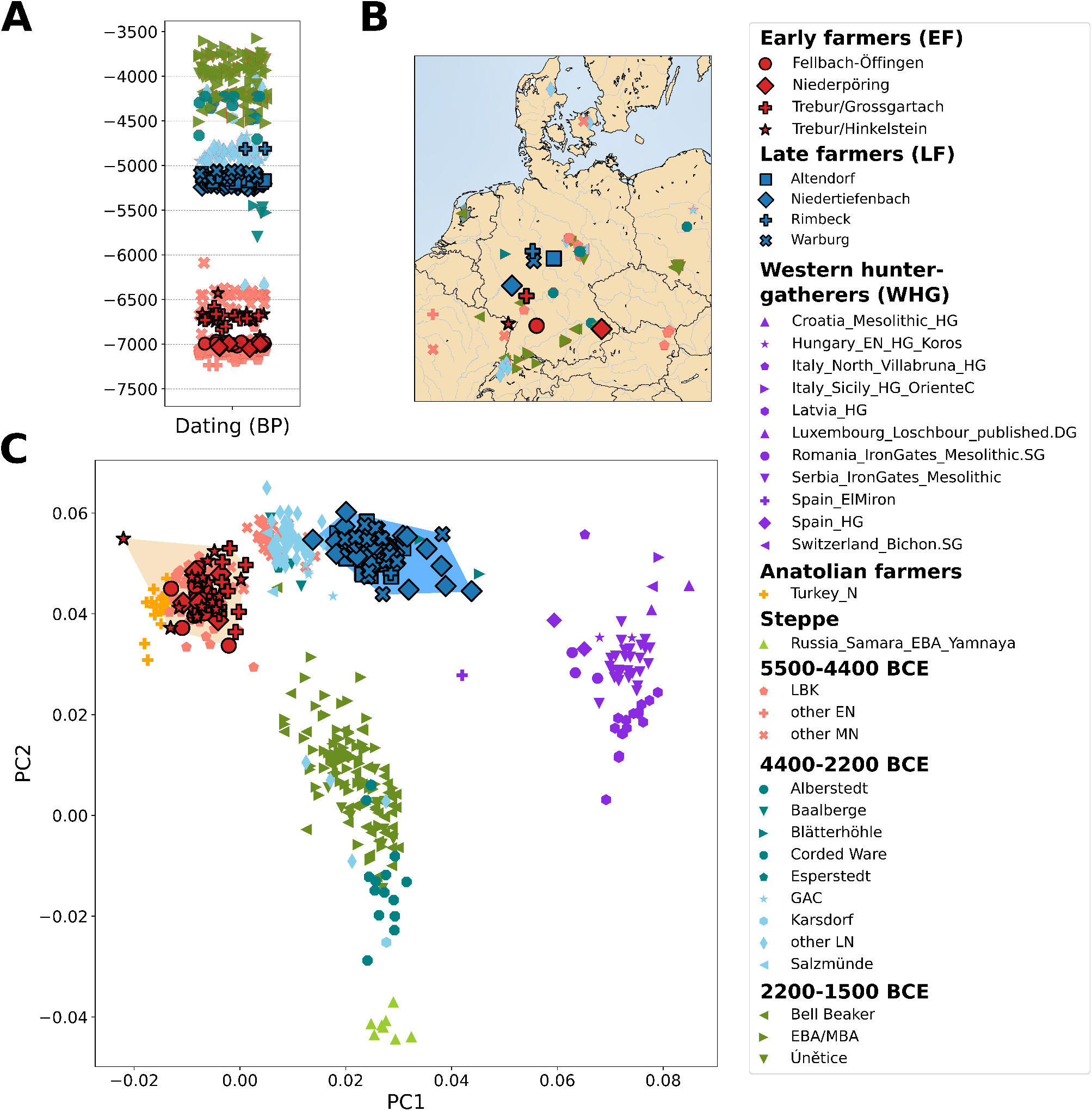
Temporal, geographic and genetic information. Timeline **(A)** and geographic map **(B)** of modern-day Germany showing locations of the early (EF, red) and late (LF, blue) Neolithic sites included in this study. Principal component analysis **(C)** of ancient individuals projected onto modern West Eurasian variation. Convex hulls highlight the space filled by EF and LF. Publicly available data from Mesolithic, Neolithic and Bronze Age populations are also included. HG = Hunter Gatherer; N = Neolithic; EN = Early Neolithic; MN = Middle Neolithic; LN = Late Neolithic; EBA = Early Bronze Age; MBA = Middle Bronze Age; GAC = Globular Amphora culture; LBK = Linear Pottery culture.

**Table 1.**
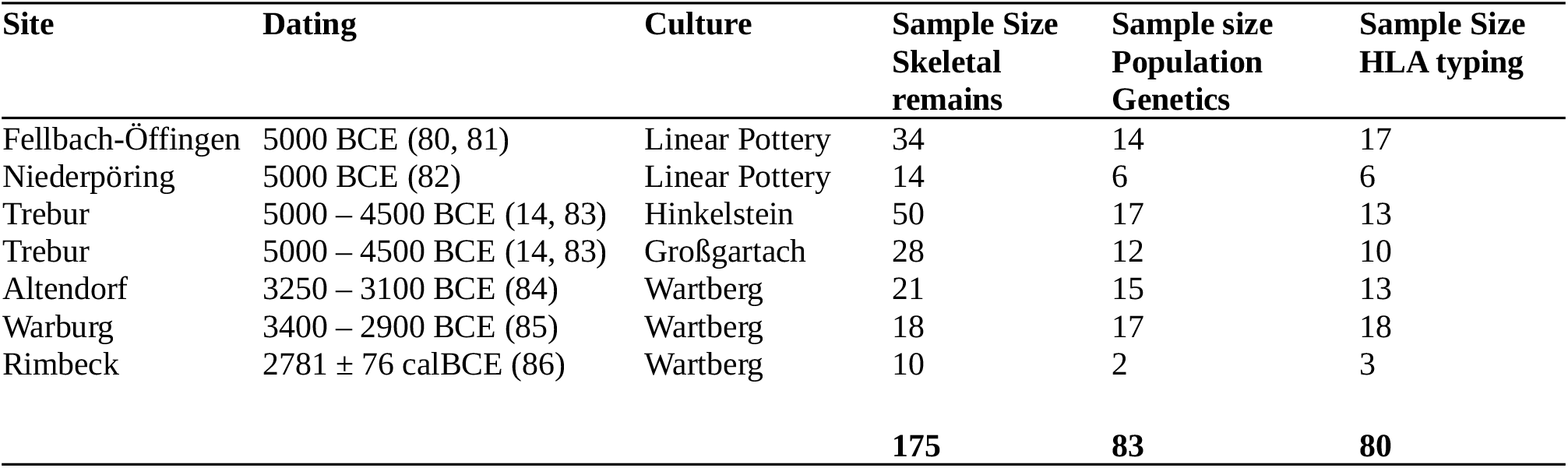
Dating, archaeological culture and sample size of the sites used in this study.

## RESULTS

In this study, we generated shotgun sequencing data (providing on average 16 million reads per library) from the human remains of 175 individuals originating from the following six sites in present-day Germany: Niederpöring, Fellbach-Öffingen, Trebur, Altendorf, Warburg and Rimbeck (Table 1; Fig. 1B; Data S1). The Trebur site contained burials assigned to the Middle Neolithic Hinkelstein and Großgartach groups (13–15).

### Metagenomic screening for pathogens

The shotgun sequencing data were screened for the presence of human blood-borne bacterial and viral pathogens. In two individuals from Niederpöring, reads of hepatitis B virus (NP560) and parvovirus B19 (NP543) were detected (Data S1). No evidence of pathogens was found in any of the other samples.

### Population genetic analyses

When mapping the shotgun sequencing data to the human genome (summary statistics available in Data S2), 83 of the 175 datasets had at least 20,000 SNPs of the 1240K panel covered and were included in the subsequent population genetic analyses (Table 1; Data S1). Principal component analysis (PCA) showed that the six populations formed two distinct groups: individuals from Niederpöring, Fellbach-Öffingen and Trebur clustered with published Early Neolithic farmers, whereas individuals from Altendorf, Warburg and Rimbeck were placed near agriculturalists from the Late Neolithic (Fig. 1C). Therefore, we will refer to the former (Niederpöring, Fellbach-Öffingen, and Trebur; n=49) as early farmers (EF) and the latter (Altendorf, Rimbeck, and Warburg; n=34) as late farmers (LF).

Unsupervised admixture analysis revealed that both EF and LF carried two major ancestry components, one maximized in WHG and the other in Anatolian Neolithic farmers (AN) (Fig. S1). EF showed higher genetic affinity with other contemporaneous groups from the LBK, Sopot and Starčevo societies (Fig. S2), while LF were more similar to WHG proxies (i.e., Loschbour Luxembourg, Bichon Switzerland, and one individual from Mont Aimé/Paris Basin). Correspondingly, two-way qpAdm models with WHG and AN as sources showed a much lower WHG component in EF (3-5%) than in LF (31-36%) (Fig S3; Data S3). We also tested three-way qpAdm models with a steppe proxy (Russia Samara EBA Yamnaya) as a third source in LF. The results suggest virtually no admixture with populations carrying the steppe-related ancestry component (Data S3). Admixture date modelling using WHG and AN as sources revealed that the WHG introgression into LF most likely occurred between the 4^th^ and 3^rd^ millennium BCE for Altendorf and Warburg (Fig. S4, Data S4). For the two Trebur subgroups, the unsupervised admixture and the PCA analyses suggested a difference in the amount of WHG ancestry. However, individual qpADM modelling did not support this (Fig. S5; Data S5).

### Kinship analysis

We explored the possibility of kinship by calculating the relatedness coefficient based on pairwise mismatch rates. Mitochondrial (mt) DNA and Y-chromosome haplogroups were also considered in the analysis (Data S1). We identified a few cases of 1^st^- or 2^nd^-degree kinship in Altendorf (n=2 relationships), Fellbach-Öffingen (n=1) and Trebur/Hinkelstein (n=6) (Fig. S6; Table S1). Consequently, one individual of each pair of related individuals was removed for the HLA frequency calculations (the one with the more complete HLA profile was kept).

### Inclusion of data from the site Niedertiefenbach

In a previous study, we described the in-depth analyses (population genetics, kinship, phenotype reconstruction, pathogen screening and HLA typing) of the Niedertiefenbach community, a Late Neolithic group associated with WBC (5). The population genetic analyses performed here showed that the Niedertiefenbach individuals clustered with LF populations, including Altendorf, Warburg and Rimbeck (Fig. 1C). In addition, the Niedertiefenbach collective displayed a large WHG ancestry component (34–58%) typically observed in the Late Neolithic (5). Radiocarbon dates (16) and cultural affiliation (WBC) (17) support the classification of Niedertiefenbach as an LF population. Therefore, the data generated from the Niedertiefenbach individuals were included in the LF group for subsequent analyses (sex-biased admixture, runs of homozygosity, HLA typing and frequency calculations).

### Sex bias in LF

From EF to LF, we observed more drastic changes in the distribution of Y-chromosome haplogroups than in the mtDNA haplogroups. For instance, LF only had one Y-chromosome macro lineage (I), whereas EF had five (Fig. S7). This finding might indicate the presence of a sex-bias during the admixture that formed LF. Therefore, we tested whether this was the case. First, we explored the statistic Q which measures relative genetic drift between the X-chromosome and autosomes. Q is expected to be 0.75 if the effective population size of males and females is equal. Deviations from this value may be suggestive of a sex-biased demography. Comparing AN and LF rendered results compatible to the expected value (Q=0.76), while the WHG–LF comparison suggested a slight deviation (Q=0.63) (Table S2). We then computed the ancestry proportions on the X-chromosome and autosomes separately and calculated the ratio of X-chromosome to autosome WHG ancestry (which we here refer to as R_X/A_). An equal admixture contribution of males and females should lead to R_X/A_=1, while deviations from this may be indicative of male or female biased admixture. We observed R_X/A_<1 in 24 out of 38 individuals (63%) that entered the analysis (mean R_X/A_=0.8; median R_X/A_=0.87; right-sided binomial test, p=0.0717; Fig. S8; Data S6). Interestingly, a few individuals (n=14; 37%) showed drastic deviations (R_X/A_<0.5) from the expected value of 1. The distributions of X-chromosome and autosome WHG ancestry in LF were significantly different (median p = 0.01, Wilcoxon signed-rank test; Fig. S9), suggestive of some WHG male-biased admixture (i.e., more WHG male ancestors).

### Runs of homozygosity

We investigated the amount of runs of homozygosity (ROH) between EF and LF. It was possible to infer ROH for 6 EF and 39 LF individuals (Fig. S10, Data S7). EF individuals presented on average shorter ROH (6cM) than LF (12cM). When we included data from published LBK sites (n=24 individuals) in the EF group to increase the sample size, the average ROH remained in the same range (5cM). We observed a statistically significant difference in average ROH between EF (including published LBK data) vs. LF (p=0.0075). ROH≥20cM were found in one individual associated with LBK and two LF individuals. We also used HapROH to estimate the effective population sizes (N_e_), which showed significantly higher values for EF (including published LBK data) (N_e_=7570, 95% CI=5105-11950) than for LF (N_e_=3371, 95% CI=2665-4384).

### HLA genotyping and analysis

For the six populations, 95 samples were subjected to in-solution DNA enrichment and sequencing of the three HLA class I (HLA-A, -B, -C) and class II loci (HLA-DPB1, -DQB1, -DRB1). Data of sufficient quality for HLA genotyping was generated for 80 unrelated individuals (Table 1). Through inclusion of data from 56 Niedertiefenbach individuals (5) to the LF group, we achieved a total of 136 HLA profiles (EF=46, LF=90) with varying levels of coverage per locus (Fig. S11G). The data was used for allele frequency calculations (genotypes available in Data S1; allele frequencies in Data S8 and Fig. S11A-F). We observed significant changes in eight HLA alleles between EF and LF (p≤0.05, Fisher’s exact test corrected for multiple testing, Fig. S12, Table S3). We additionally found major changes (>=10% frequency difference) in 17 alleles between the Neolithic groups (either EF or LF) and a representative sample of modern Germans (18) (p≤0.05, Fisher’s exact test corrected for multiple testing, Fig. S12, Table S3). The largest frequency changes (≥20%) in both comparisons affected mostly the HLA class II loci. Furthermore, we noted the co-occurrence of some HLA class II alleles, indicating possible haplotypes: DRB1*13:01-DQB1*06:03 with 19% frequency in EF and DRB1*08:01-DQB1*04:02 with 20% frequency in LF. Out of the 20 most common HLA alleles in modern Germans (≥10%), 7 alleles were not observed in our Neolithic samples and 6 were present only at low frequencies (<5%) (Table S4).

To compare the HLA diversity between the two Neolithic groups and modern Europeans (Germans (18) and five populations with European ancestry from the 1000 Genomes dataset (19, 20)), we applied the Shannon diversity (21) index to quantify the HLA allele diversity within each population. The diversity was consistently and significantly lower in EF compared to LF in all loci except for HLA-A and HLA-B (Fig. 2, Data S9). Similarly, both EF and LF had lower diversities relative to modern populations except for HLA-A and HLA-C. The most drastic difference was observed in HLA-DQB1 that showed a remarkably low diversity in both Neolithic groups, owing to the dominance of a few HLA-DQB1 alleles during that period. The two most common HLA-DQB1 alleles in EF and LF reached cumulative frequencies of 80% and 65%, respectively, while a more even distribution of frequencies was observed in modern Germans (Fig. S11).

**Fig. 2.**
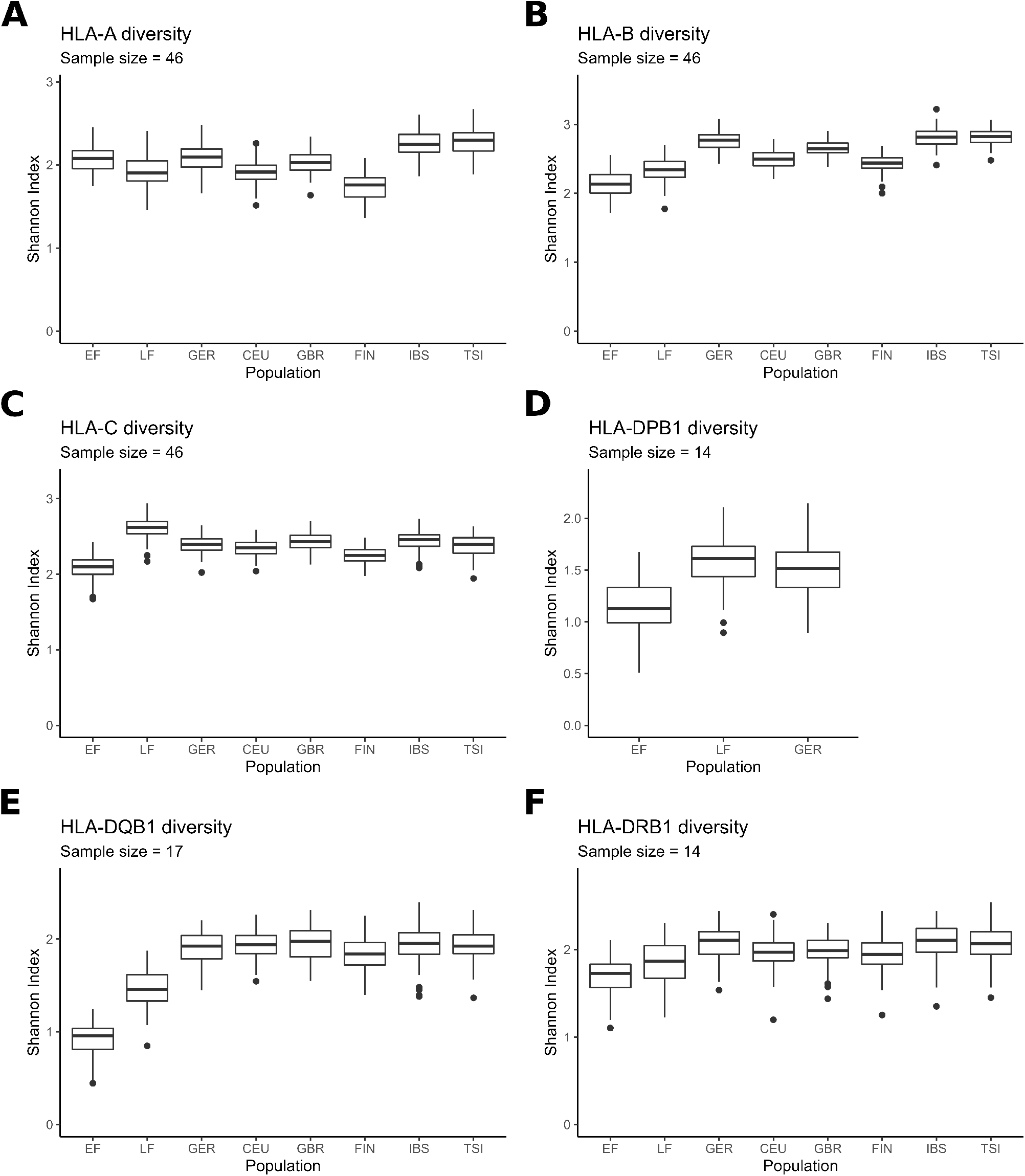
HLA diversity in the Neolithic populations presented in this study compared to modern populations. HLA diversity was measured by the Shannon index (H’) for the loci A **(A)**, B **(B)**, C **(C)**, DPB1 **(D)**, DQB1 **(E)** and DRB1 **(F)**. Boxplots represent the distribution of the H’ values of 100 samples taken from each population with sample sizes indicated below headers. EF = early farmers; LF = late farmers; GER = modern Germans; CEU = Central Europeans; GBR = British; FIN = Finnish; IBS = Iberians from Spain; TSI = Tuscans from Italy.

## Discussion

Here, we generated genome-wide data for 83 individuals from six archaeological sites in present-day Germany that cover the Early Neolithic (Fellbach-Öffingen, Niederpöring, and Trebur) and Late Neolithic (Altendorf, Rimbeck, and Warburg) (Table 1). Given the genetic commonality among the Early Neolithic populations (Fig. 1), we referred to them here under the term early farmers (EF; n=49). Correspondingly, the three Late/Final Neolithic WBC communities were designated as late farmers (LF, n=34).

Our analyses showed that EF, like individuals from other published LBK sites in Germany (3, 22, 23), closely resembled Anatolian farmers (95-100% Anatolian ancestry component). In addition, they carried mtDNA and Y-chromosome lineages characteristic of early farming populations (2, 24) (Fig. S7). The genetic continuity throughout the LBK indicates long-lasting intracultural mating practices. However, close-kin mating was likely prevented as the analysis of ROH and N_e_ suggested a large group size and a wide partner exchange network as reported previously (11, 25).

The LF studied here were characterized by a high WHG ancestry proportion (31-36%), the influx of which may also have led to changes in mtDNA and Y-chromosome lineages (Fig. S7). Our analyses in LF individuals suggested a statistically significant male-bias from WHG during their admixture with early farmers (Figs. S8 and S9). LF presented more and longer ROH than EF. One explanation for this finding could be recent admixture with WHG introducing longer ROH. Another scenario could be the mating of relatives. However, the latter is not supported by our data, as mostly unrelated individuals were detected in the four collective burials studied here (Altendorf, Niedertiefenbach, Rimbeck, Warburg) or in the gallery grave of Niedertiefenbach (5). During the EF to LF transition, farming communities appear to have changed from closed to more permeable societies that were willing and able to integrate WHG, a process that was further accompanied by the diversification and regionalization of archaeologically defined groups.

Our genetic dating of the admixture event between WHG and EF confirmed our previous results that WBC (LF) most likely emerged from a Michelsberg context (MC; 4400–3500 BCE) (5). There is evidence suggesting that the MC farmers were particularly mobile. For example, some MC groups used flint from non-local quarries, indicating that they were engaged in long-distance barter. In addition, they practised forest pasture management which can be interpreted as transhumance (26–28). This mobility may have led MC people to increasingly engage with WHG, contributing to the admixture of both groups. It is possible that the cultural characteristics of the admixed WBC groups were influenced in part by the relative contributions of each ancestral population (i.e., WHG and LBK). The admixture event represented a profound transformation with long lasting effects on demography, gene pool and culture in Europe.

Next, we investigated whether EF and LF differed in their HLA variation. For eight HLA alleles, we observed significant frequency differences (Table S3). As EF and LF varied in their proportion of WHG ancestry, the most plausible explanation for the considerable frequency shifts is admixture with WHG rather than selection on each allele. This is supported by our results showing an increased HLA diversity in LF compared to EF (Fig. 2). However, the hypothesis remains to be further tested once true HLA calls become available for WHG. Interestingly, the major frequency changes (>20%) between EF and LF mainly affect HLA class II alleles (Table S3). A recent SNP-based study (29) has shown that the HLA region, especially the DQB1 locus (HLA class II), is enriched for WHG ancestry in Neolithic individuals as a result of adaptive admixture. This process has been shown to be relevant within the HLA context, potentially driving specific alleles towards higher frequencies (30, 31). The HLA allele spectrum of WHG may have differed in part from that of EF because they had been adapted to the European environment and a hunting-gathering lifestyle. Taking these findings into account, we hypothesise that the increase in frequency of specific HLA class II alleles (such as DQB1*04:02) observed in LF could have been affected by adaptive admixture.

Interestingly, the alleles with high frequencies in EF, DRB1*13:01 (27%) and DQB1*06:03 (31%), whose co-occurrence indicates a haplotype, have been shown to be protective against viral hepatitis A (HAV) and B (HBV) infections today (32–34). In LF, these alleles were much rarer (6% and 10%, respectively); instead, LF had high frequencies of DRB1*08:01 (27%) and DQB1*04:02 (34%), which may form another haplotype that is also protective against HBV (35). It appears that the increase/decrease in the two potential haplotypes is proportional, indicating that the functional protective effect against HBV was maintained. aDNA studies have shown that HBV was already endemic in WHG and Neolithic populations (36, 37), albeit in form of phylogenetically distinct strains. The occurrence of frequent protective HLA class II alleles (though different ones) suggests that the virus may have been a strong selective pressure in both WHG and farmers.

An additional factor contributing to the HLA shifts between EF and LF may be changing pathogen landscapes throughout the Neolithic. It has been hypothesized that the adoption of the Neolithic lifestyle (e.g., sedentary groups living closely with domesticated animals) was associated with an increase in infectious diseases and epidemics (38). However, archaeological and aDNA studies have so far not provided evidence for large-scale epidemics, only sporadic infections caused by a very limited number of pathogens (36, 37, 39, 40). Based on current data, this low pathogen load did probably not change from the Early to the Late Neolithic.

With regard to the HLA allele repertoire, it is noteworthy that EF and LF had a relatively low diversity compared to modern populations. This means that a few alleles were observed at exceptionally high frequencies (>20%) (Fig. S11, Data S8). Theory predicts that the presence of such common HLA alleles over extended periods of time should increase the probability that pathogens evolve evasion mutations to reduce the likelihood of their recognition by the immune system (41). The maintenance of the frequent alleles over two millennia might therefore support the observation that the low pathogen threat and load likely remained the same throughout the Neolithic.

When comparing the two Neolithic groups with modern Germans, we observed significant changes in the frequencies of 17 HLA alleles (Table S3). Fourteen alleles showed a significant decrease in frequency towards the present and three alleles followed the reverse trend. The most drastic changes affected HLA class II alleles. However, noteworthy are also the HLA class I alleles HLA*B:27:05 and HLA-B*51:01 which are strongly associated with inflammatory diseases (ankylosing spondylitis and Behçet’s disease, respectively) (42–45). Their high frequency in the Neolithic has been previously observed (5, 11).

Surprisingly, seven HLA alleles present at high frequencies (≥10%) in today’s Germany were virtually absent in EF and LF (Table S4). This finding suggests that these common HLA alleles were likely introduced after the Neolithic period. Their increased frequency can be due to admixture processes (e.g., with groups carrying the steppe-related ancestry), pathogen-driven selection (e.g., negative frequency-dependent selection, or directional selection by novel pathogens) or a combination of both. To address these questions, more palaeogenomic studies with true HLA calls as well as a better characterization of the pathogen landscape are needed for populations before and after the Neolithic.

## METHODS

### Sampling

In total, we sampled 185 human remains from six sites within Germany ranging from the Early to the Late Neolithic (Figs. 1A-B, Table 1).

### DNA extraction and library preparation

DNA was extracted from teeth and/or bones of all individuals and converted into partial Uracil-DNA Glycosylase (UDG) libraries (46) following established laboratory guidelines for aDNA work (47). Shotgun sequencing was performed on the Illumina HiSeq 6000 (2×100) platform of the Institute of Clinical Molecular Biology (IKMB) in Kiel. Additionally, UDG-treated libraries were enriched for the HLA region applying a custom bait capture (48). The targeted capture was conducted on 95 samples, of which 80 samples were successfully enriched to be analyzed.

### Metagenomic screening

The sequencing reads were screened for the presence of pathogens following an in-house pipeline (49, 50) using MALT (51) v0.4.1 with a semi-global alignment mode and a minimum percent identity of 90% to align the samples against a database of 27,730 bacterial and 10,543 viral complete genomes (52, 53).

### Mapping

The removal of adapter sequences as well as the merging of paired-end reads were performed with ClipAndMerge (54) v1.7.7. Mapping to both the human genome (build hg19) and human mitochondrial genome references was done with BWA (55) v0.7.15 using reduced mapping stringency settings (flag -n 0.01) to account for mismatches expected in aDNA. Duplicates were removed with DeDup (54) v0.12.1.

### Contamination estimation and genetic sex determination

To evaluate the authenticity of samples as ancient, we assessed terminal damage of reads by calculating the frequency of C to T substitutions with DamageProfiler (56) v1.1. After validation, the first two positions from the 5′ and 3′-ends of the reads were removed with bamUtil (57) v1.0.15. Mitochondrial DNA contamination was estimated by analysing sequence deamination patterns and fragment length distributions with Schmutzi (58) v1.5.5.5. Additionally, contamination in male samples was measured by assessing X chromosome heterozygosity with ANGSD (59) v0.935. Samples that showed more than 5% mtDNA or X chromosome contamination were excluded from further analysis. In cases where contamination estimation with Schmutzi was not possible, the placement of the individuals in the PCA plot was additionally used to further assess if the samples should be excluded. Sex was genetically determined by considering the ratio of sequences aligning to the X chromosome and autosomes (60). Only samples with more than 1,000 reads were considered for sex determination.

### Genotyping

SequenceTools (https://github.com/stschiff/sequenceTools) v1.2.2 was used to generate pseudo-haploid genotypes on 1,233,013 SNP positions (4, 22, 61). Samples with fewer than 20,000 genotyped SNPs were excluded from the analysis.

### Mitochondrial and Y chromosome haplogroups

Mitochondrial haplogroups were determined with HaploGrep2 (62) and Y haplogroups with yHaplo (63). A mapping and base quality threshold of 20 was used. For Y haplogroups, the presence of at least 10 derived alleles was used as a threshold to make a call.

### Principal component analysis

The genotyped samples in this study were merged with the Allen Ancient DNA Resource (AADR) reference panel (v50.0.p1) containing previously published genotypes of 10,342 ancient and modern individuals (64). The PCA was performed with *smartpca (65)* from the EIGENSOFT package and with the “lsqproject” option. The calculation of principal components was based on a subset of 66 modern populations from West-Eurasia (Human Origins samples in the AADR dataset), while the remaining individuals from the merged dataset were projected into that space.

### Outgroup f3 statistics

Shared genetic drift was calculated with the program *qp3Pop* from the Admixtools package (66) in the format f_3_ (sample population; test population, Mbuti), where “sample population” refers to the 6 new populations described in this study and “test population” refers to published ancient groups available in the 1240K SNP panel.

### Admixture analyses

The merged genotype data was pruned in PLINK (67) v1.90b6.21, with an r^2^ threshold of 0.4, a window size of 200 and a step size of 25 (command “indep-pairwise 200 25 0.4”). An unsupervised ADMIXTURE (68) v1.3.0 analysis was performed on the resulting dataset, using a range of 4 to 12 components (*K*) with 100 bootstraps each. Cross-validation error was calculated for each model to identify the best *K* component. Two- and three-way admixture models were tested with *qpAdm* from Admixtools (66). DATES was used to estimate time of admixture, with the parameters ‘binsize’: 0.001, ‘maxdis’: 1.0, ‘seed’: 77, ‘jackknife’: YES, ‘qbin’: 10, ‘runfit’: YES, ‘afffit’: YES, ‘lovalfit’: 0.45, ‘minparentcount’: 1 (69). A generation time of 29 years was used to calculate the admixture calendar years (70). The potential sources and outgroup populations for the *qpAdm* and DATES analyses are listed in Data S3, S4 and S5.

### Sex-biased admixture in late farmers

As a first measure to assess sex-biased admixture in LF (Altendorf, Warburg, Rimbeck and previously published Niedertiefenbach), we compared genetic differentiation on the X chromosome (F_st_X) and autosomes (F_st_A) between LF and the two source populations: WHG and Anatolian Neolithic farmers (AN). For this, we used the SNPs belonging to the 1240K panel and filtered out positions with r^2^>0.4 (plink command “indep-pairwise 200 25 0.4”). SNPs in the the pseudoautosomal regions of the X chromosome were also removed. After filtering, 560,930 autosomal and 5,004 X chromosomal SNPs remained. Individuals with fewer than 1,000 SNPs covered on the X chromosome were removed from the analysis. We then used the obtained Weir and Cockerham weighted F_st_ values to calculate the statistic Q (71–73). This statistic measures relative genetic drift between the X chromosome and autosomes and is calculated as

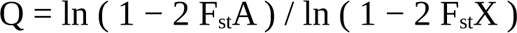

As Q can be influenced by factors other than sex-biased admixture (71–73), we also computed the amount of WHG ancestry on the X chromosome versus autosomes using supervised ADMIXTURE analyses with WHG and AN as sources. As the number of SNPs available for the analyses on the X chromosome is low (n=5,004) compared to autosomes (n=560,930), we resampled autosomal SNPs 1,000 times relative to the number of SNPs available for the X. For each individual, we calculated the ratio of WHG ancestry on the X chromosome versus autosomes. We used the Wilcoxon signed-rank test to assess significant differences between the means of X and autosomal WHG ancestry (71).

### Kinship analysis and runs of homozygosity (ROH)

To estimate kinship, we used the method described in Fowler et al. (74). Shortly, for each pair of individuals we calculated pairwise allelic mismatch rates in autosomal sites of the 1240K panel. We then computed relatedness coefficients *r* for each pair using the formula

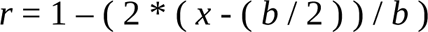

where *x* is the mismatch rate of the pair of individuals and b the expected mismatch rate for two unrelated individuals from the same population. To calculate the constant *b*, we first merged data from our six populations (n=83 individuals) with published data from 15 Neolithic populations located in present-day Germany (n=155 individuals). Then, we calculate pairwise mismatch rates for all combinations of two individuals from the merged dataset (28,203 comparisons) and used bootstrapping to calculate 95% confidence intervals. We filtered out pairwise comparisons with fewer than 100K overlapping SNPs (7,101 comparisons remained after filtering) and calculated *b* as the median mismatch rate of the filtered dataset (*b*=0.2593), a value similar to that obtained by Fowler et al. (74) (0.2504) using Neolithic individuals from England. We then applied our obtained value of *b* in the formula described above to calculate the relatedness coefficient for each pair of individuals. Relationship degrees were annotated using the same cutoffs as in Fowler et al. (74). Pairwise comparisons with fewer than 2500 overlapping SNPs or with a large confidence interval leading to annotation of more than 2 possible degrees of kinship were not considered. Mitochondrial DNA and Y chromosome haplogroups, when available, were also considered in assessing kinship. We screened for ROH using HapROH (25) with the default parameters. Only samples with more than 400,000 SNPs genotyped from the 1240K panel were included. The results were merged with previously published ROH estimations (25). Due to the small sample size of EF, seven published populations were added for calculating the average sum of ROH (Data S7). The ROH results were then used to infer the effective population size (N_e_) also with HapROH, using the default parameters. A Mann-Whitney U test was performed with the python3 module scipy v1.9.1 to test for significant differences in the average sum of ROH between groups.

### HLA genotyping and frequency calculations

Genotyping of the HLA alleles was performed for the three class I (HLA-A, -B and -C) and three class II (HLA-DPB1, -DQB1 and -DRB1) loci using a combination of OptiType (75) and the TARGT pipeline (Targeted Analysis of sequencing Reads for GenoTyping) (76), which was designed for the analysis of low-coverage sequences such as ancient DNA data. To ensure a higher reliability of the results, the manual genotyping was done by two independent scientists and HLA class I calls were additionally verified using OptiType. Only alleles consistently called by both methods were included. All analyses were done at two-field HLA allele resolution. For the allele frequency calculations, we grouped the populations according to their dates, cultural affiliation and population structure as EF (Niederpöring, Fellbach-Öffingen, and Trebur) and LF (Altendorf, Rimbeck, Warburg and Niedertiefenbach). We excluded from the allele frequency calculations seven individuals from seven kinship clusters containing 1^st^ degree relationships (Altendorf=2, Fellbach-Öffingen=1, Trebur/Hinkelstein=4). For Niedertiefenbach, we included 56 HLA profiles in the LF group, 33 of which were generated as part of this study and 23 of which were previously published (5). This addition increased our data set to 47 individuals for EF and 90 individuals for LF (Data S1). For 22 individuals, the targeted HLA capture was successful, but no shotgun data of sufficient quality (see Methods) was available for them to allow population genetic analysis. However, both the archaeological context and aDNA damage plots, which showed distinct deamination patterns, demonstrated the ancient origin of the samples used (Data S2) and thus these were kept for the HLA frequency calculations. For comparison with modern Germans (n=3,456,066 (18)), data from the Allele Frequency Net Database (77) were accessed. We used Fisher’s exact test to assess whether the observed allele frequencies between groups were significantly different. The p-values were corrected for multiple testing with the two-stage Benjamini and Hochberg procedure using the python3 module statsmodels v0.13.5. Frequencies of the possible haplotypes DRB1*13:01-DQB1*06:03 and DRB1*08:01-DQB1*04:02 were calculated using the expectation-maximization algorithm implemented in the Arlequin v3.5 software (78).

Shannon’s diversity index (H’) was calculated by using the *diversity* function from the *vegan* R package to measure the genetic diversity of HLA alleles in Neolithic and modern populations. Five populations with European ancestry from the 1,000 Genomes Project (19, 20), namely British from England and Scotland (GBR), Finnish in Finland (FIN), Iberian populations in Spain (IBS), Toscani in Italy (TSI) and Utah residents (CEPH) with Northern and Western European ancestry (CEU) were included in the analysis to obtain a better estimation of the modern HLA diversity. We used a down-sampling approach to control for differences in sample sizes between ancient and modern populations, since Shannon’s diversity index uses proportions of alleles which can be affected by sample sizes. Specifically, for each locus, we first identified the population with the smallest sample size (*n*), which was always EF, and calculated allele frequencies for each population. Then, we generated 100 random samples for each population with size *n* based on the allele frequencies of that population and calculated Shannon’s diversity index. We compared the distribution of Shannon’s diversity index values between populations using the Kruskal-Wallis and Dunn’s tests.

## Funding

This study was funded by the Deutsche Forschungsgemeinschaft (DFG, German Research Foundation) CRC 1266 project ID 290391021, Cluster of Excellence ROOTS (EXC 2150 ID 390870439) and RTG 2501 ID 400993799. TLL was funded by the DFG – 437857095.

## Author contributions

B.K.-K. developed the idea for this study. S. Sch.-L., J.W., C.B., M.F., I.G., K.Sch., J.P. and G.G. assembled archaeological material. B.K.-K. was responsible for generating ancient DNA data. N.A.d.S. performed population genomic analysis. M.H. generated HLA calls and performed pathogen screening. N.A.d.S., O.Ö., Y-R.Ch., T.L.L. analysed the HLA data. N.A.d.S., O.Ö., M.H., D.K., S. Sch.-L., J.W., C.B., M.F., I.G., K.Sch., J.P., G.G., Ch.R., J.M., T.L.L., A.N., B.K.-K. interpreted the findings. N.A.d.S., A.N., B.K.-K. wrote the manuscript with major contributions from T.L.L., O.Ö. as well as input from all other authors.

## Competing interests

The authors declare that they have no competing interests.

## Data and materials availability

All data needed to evaluate the conclusions in the paper are present in the paper and/or the Supplementary Materials. Aligned sequencing reads for samples reported in this study are available from European Nucleotide Archive (ENA), accession no: PRJEB53796.

## Ethics declarations

This study was carried out following the principles for ethical DNA research on human remains as described in (79).

## Notes

### Competing Interest Statement

The authors have declared no competing interest.

