## Supplementary Materials for "Admixture as a source for HLA variation in Neolithic European farming communities"

**This PDF file includes:**

Figs. S1 to S12
Tables S1 to S4

**Other Supplementary Materials for this manuscript include the following:**

Data S1 to S9

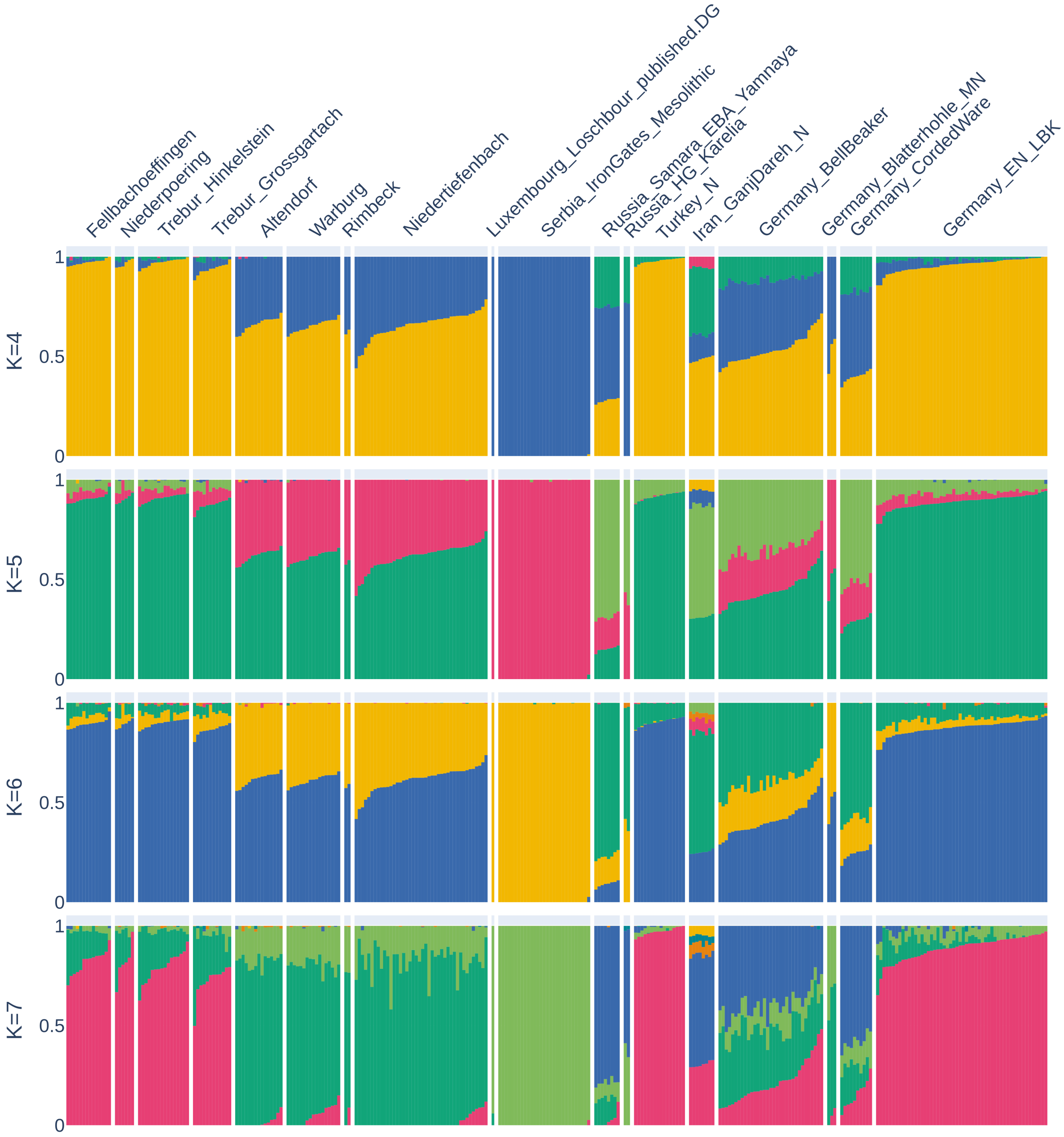


**Figure S1. Results from unsupervised ADMIXTURE.** The analysis was performed using three to twelve components for 306 selected ancient and modern populations/individuals (only 11 are plotted). The four components with lowest cross-validation values are shown. Each K was run with 100 bootstraps. HG = Hunter Gatherer; N = Neolithic; EN = Early Neolithic; MN = Middle Neolithic; EBA = Early Bronze Age; LBK = Linear Pottery culture.

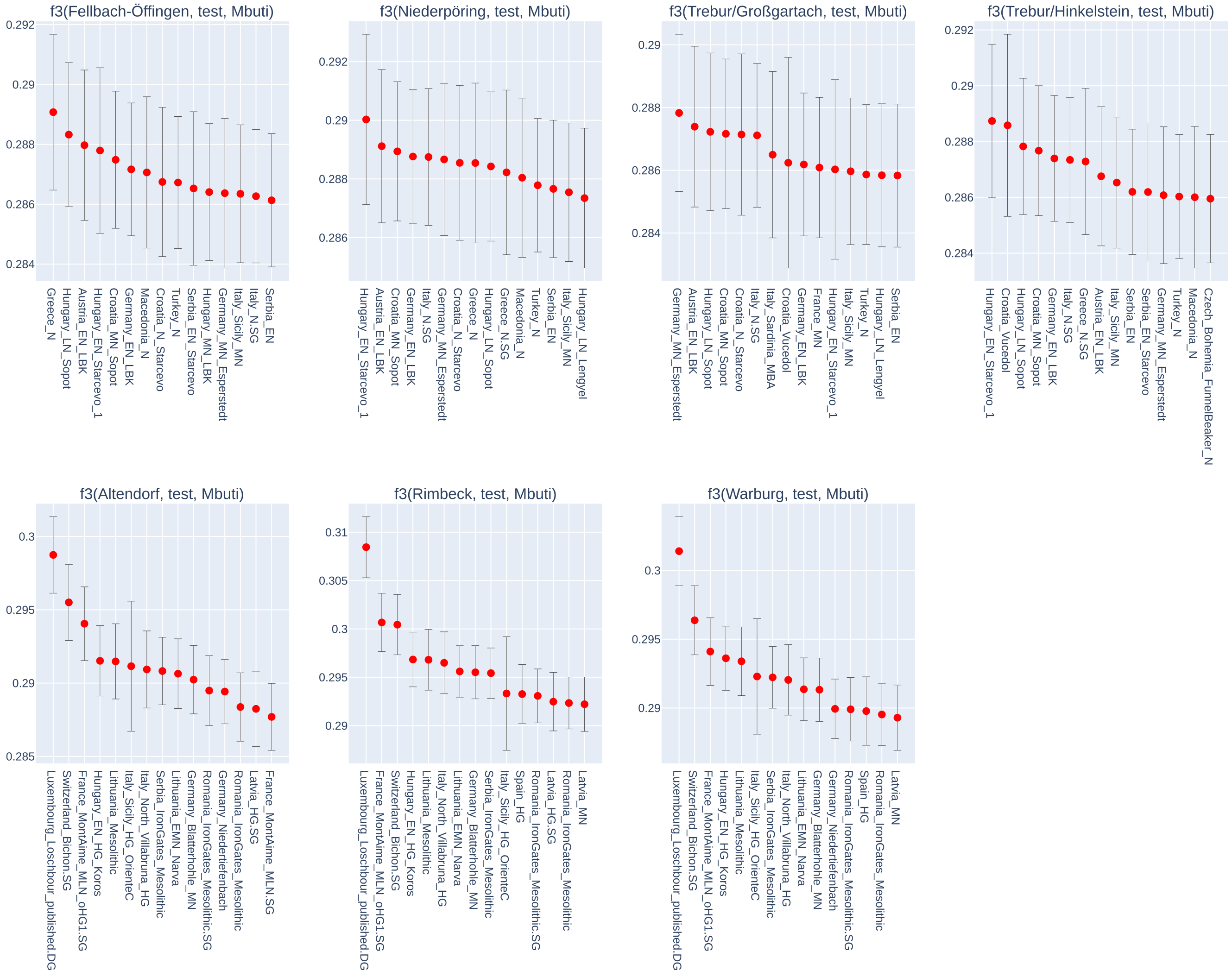


**Figure S2. f3-outgroup statistics.** The tests were performed in the format f3(Pop; Test, Mbuti) to assess the amount of shared genetic drift between each of the populations presented in this study (Pop) and other ancient populations (Test). The 15 populations with highest f3 score are shown. HG = Hunter Gatherer; N = Neolithic; EN = Early Neolithic; MN = Middle Neolithic; LN = Late Neolithic; MBA = Middle Bronze Age; LBK = Linear Pottery culture.

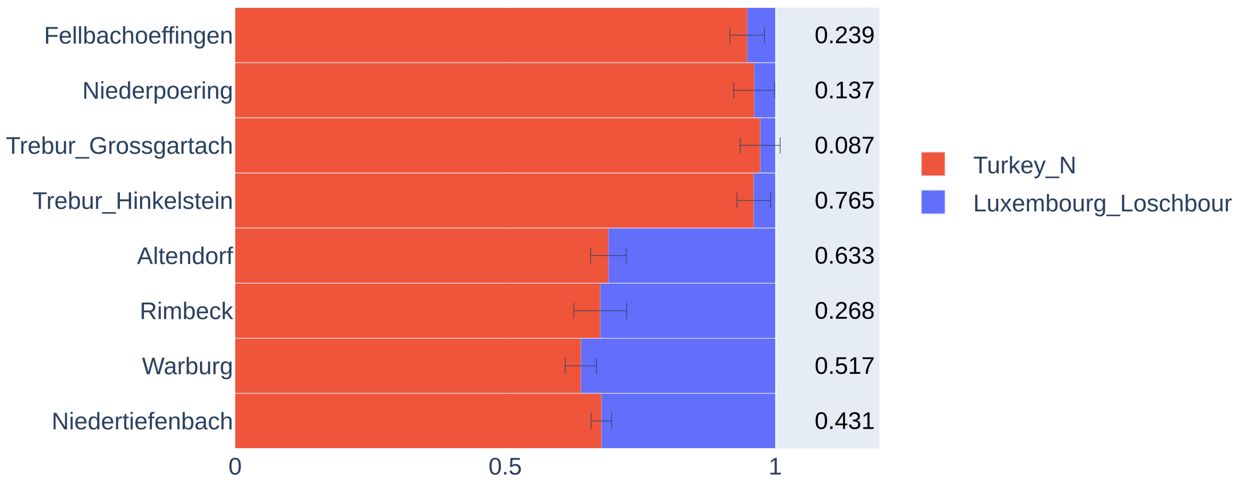


**Figure S3. qpAdm-based admixture models using two sources (Turkey_N and Luxembourg_Loschbour)**. The p-values of the feasible models are shown on the right.

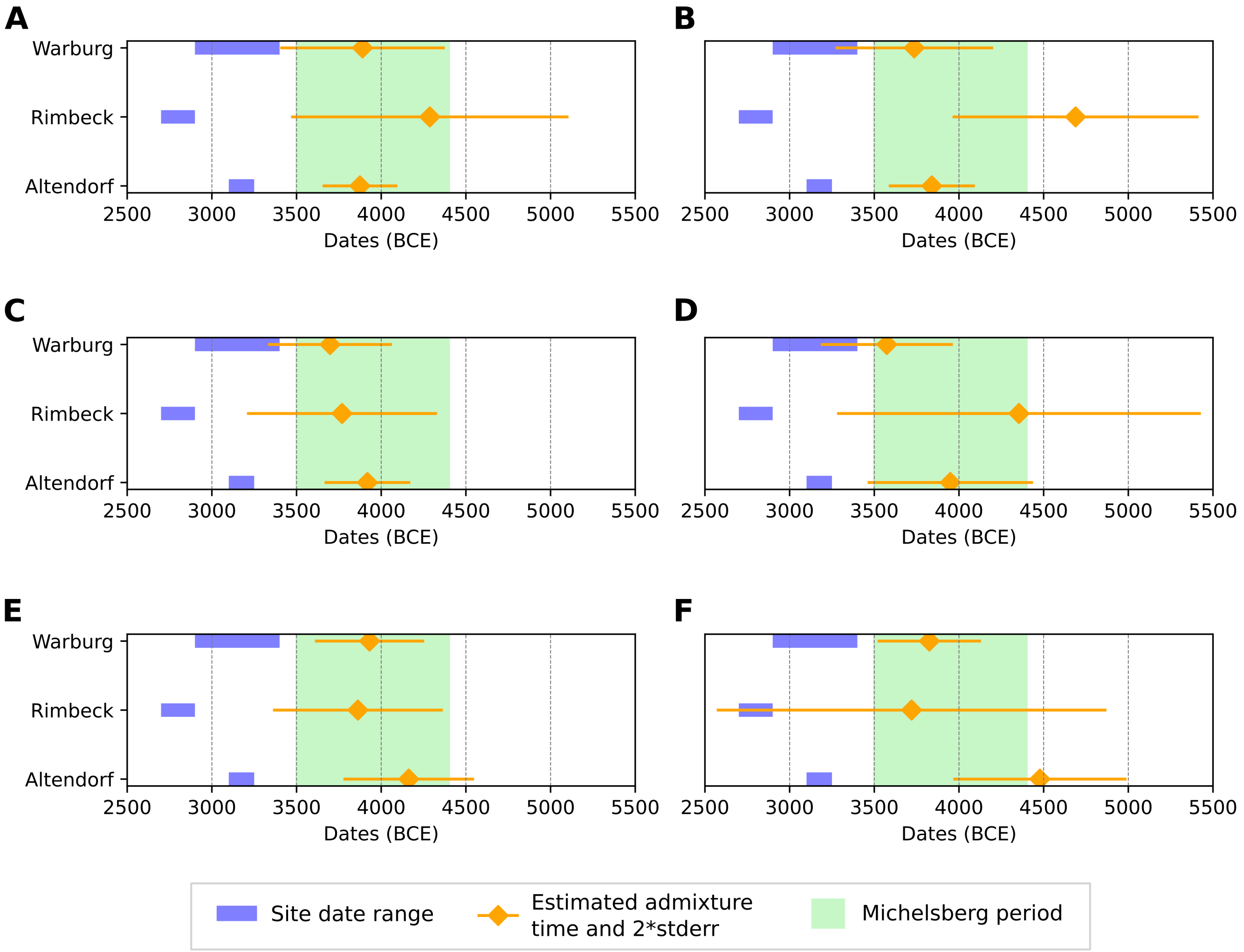


**Figure S4. DATES results.** Modelling of admixture dates using different potential source groups for the late farmer (LF) populations presented in this study: **(A)** Germany_EN_LBK and Luxembourg_Loschbour, **(B)** Turkey_N and Luxembourg_Loschbour, **(C)** Germany_EN_LBK and Romania_IronGates_Mesolithic, **(D)** Turkey_N and Romania_IronGates_Mesolithic, **(E)** Germany_EN_LBK and Switzerland_Bichon.SG , **(F)** Turkey_N and Switzerland_Bichon.SG. A generation time of 29 years was used to calculate the approximate calendar date.

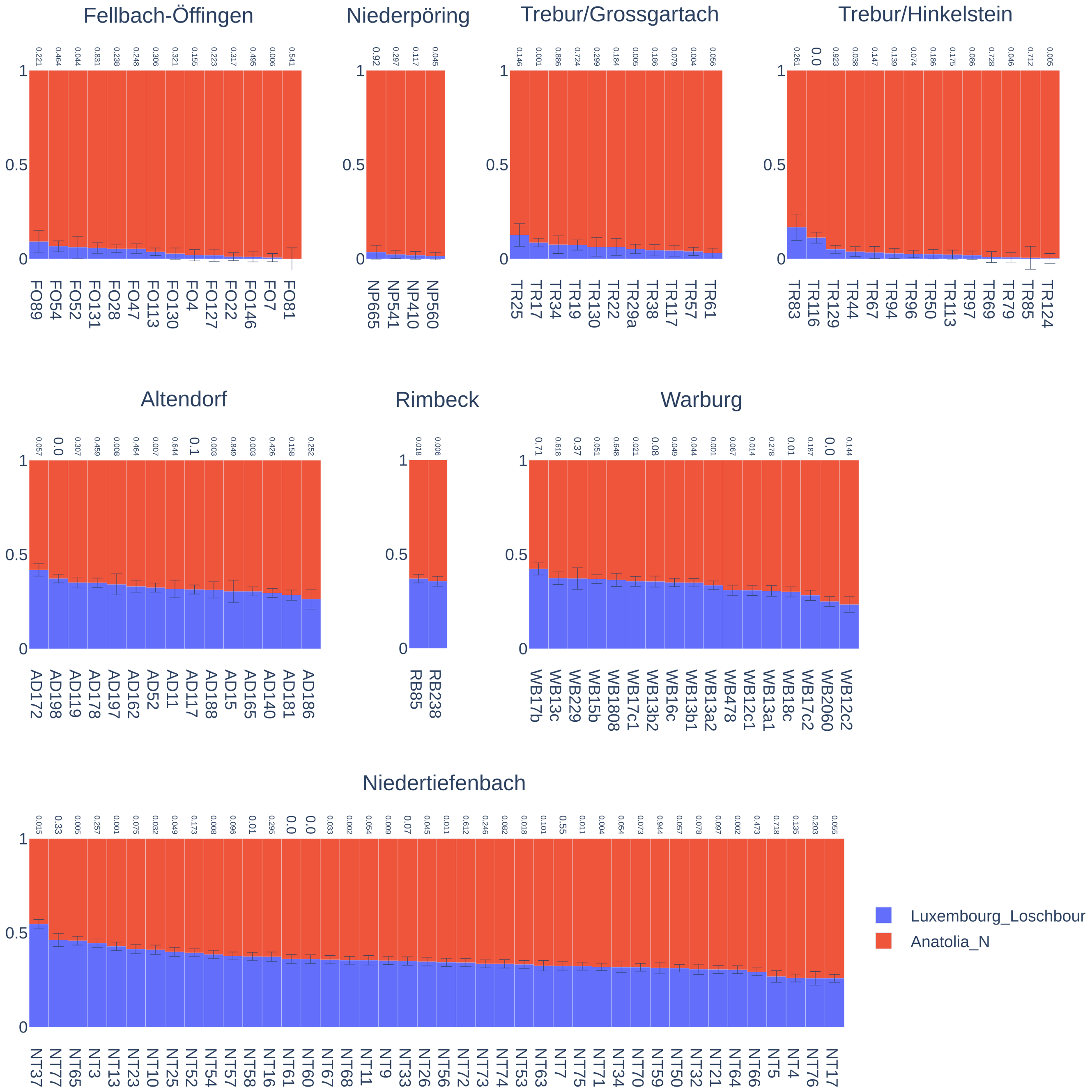


**Figure S5. Individual qpAdm-based admixture models using two sources (Turkey_N and Luxembourg_Loschbour).** Only individuals with feasible models are shown. The values on top of the stacked bars refer to the p-value of each individual model.

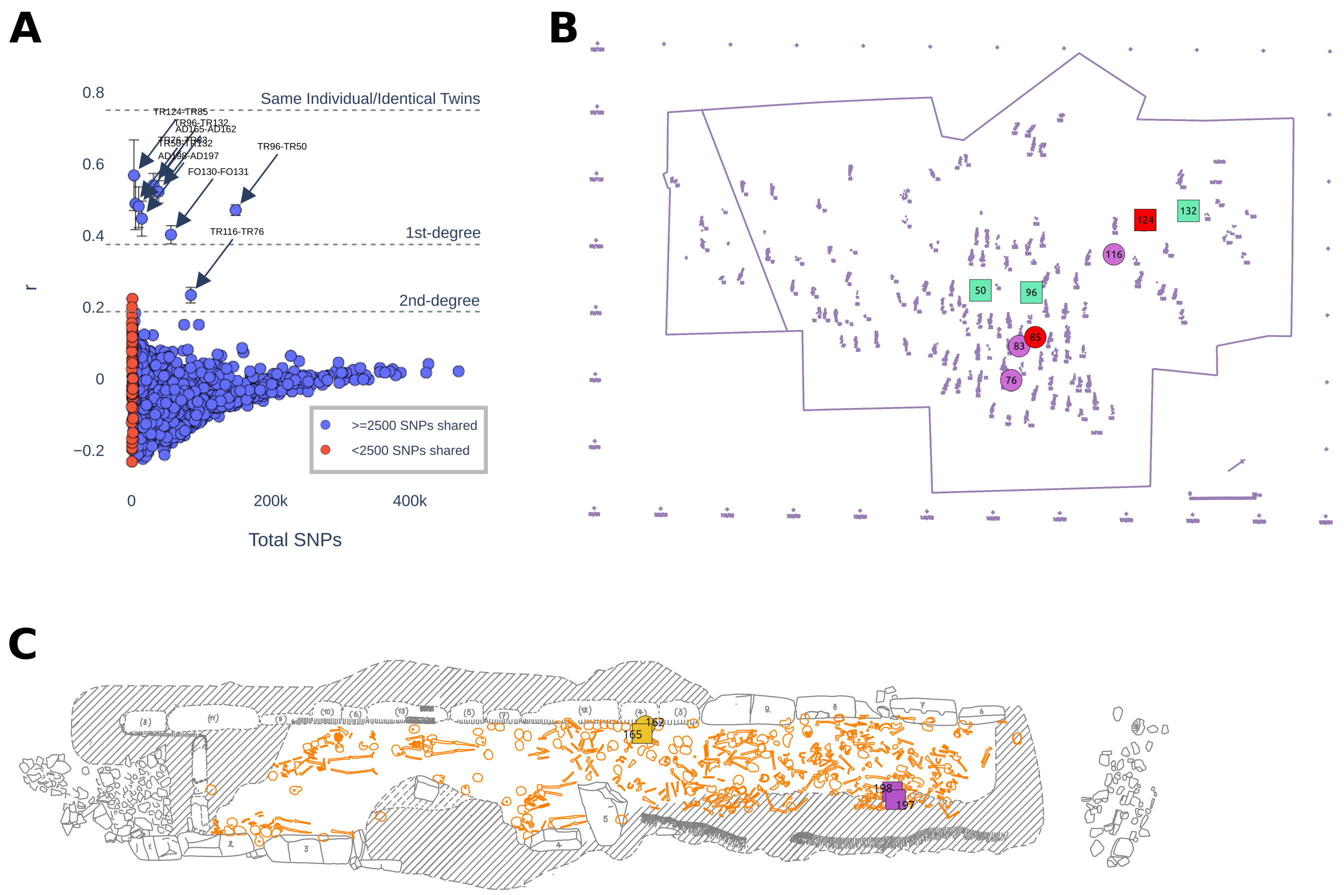


**Figure S6. Results of kinship analysis.** The first panel **(A)** displays the estimated relatedness coefficient (r) based on pairwise mismatch rate. Each marker represents one pairwise comparison. The x-axis shows the number of overlapping SNPs between each pair. The arrows point to the comparisons that correspond to the observed 1$^{st}$ and 2$^{nd}$ degree relationships described in the Results. Grave plan of the sites of Trebur **(B)** and Altendorf **(C)** are also shown, with the respective related individuals highlighted in the same colour.

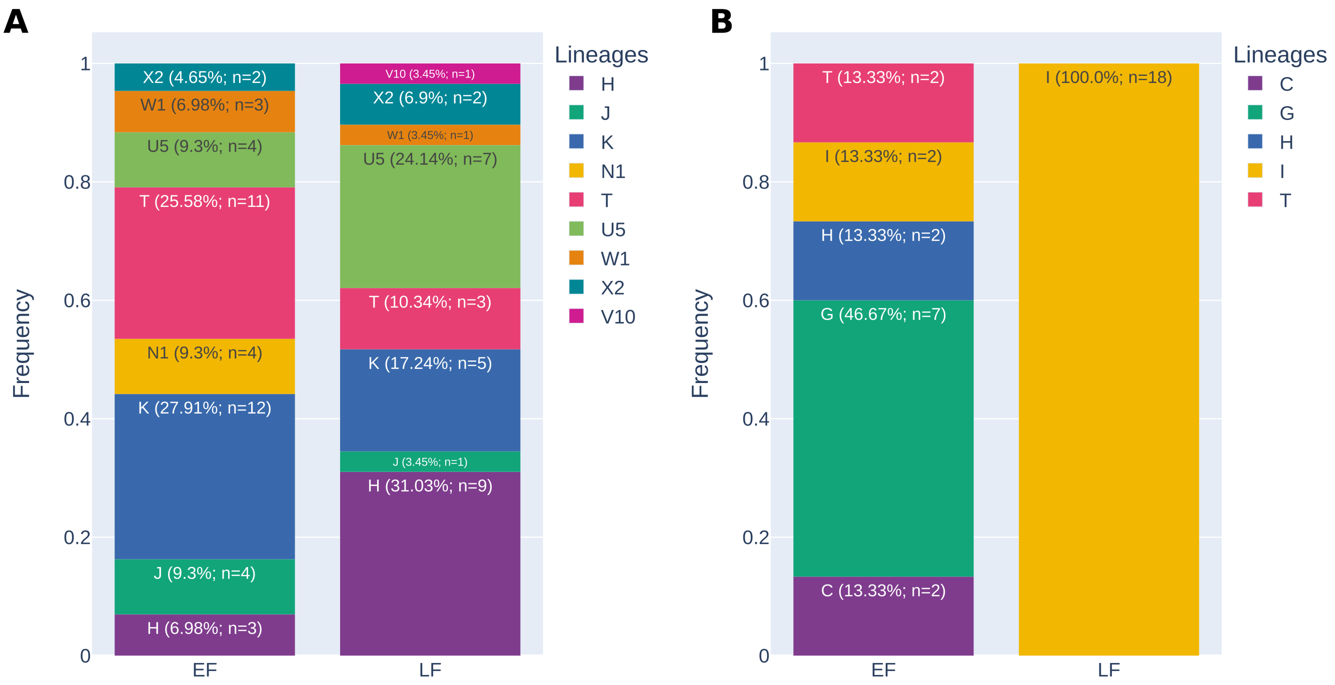


**Figure S7. Haplogroup estimation.** Distribution of mitochondrial **(A)** and Y-chromosome **(B)** macro lineages in EF and LF.

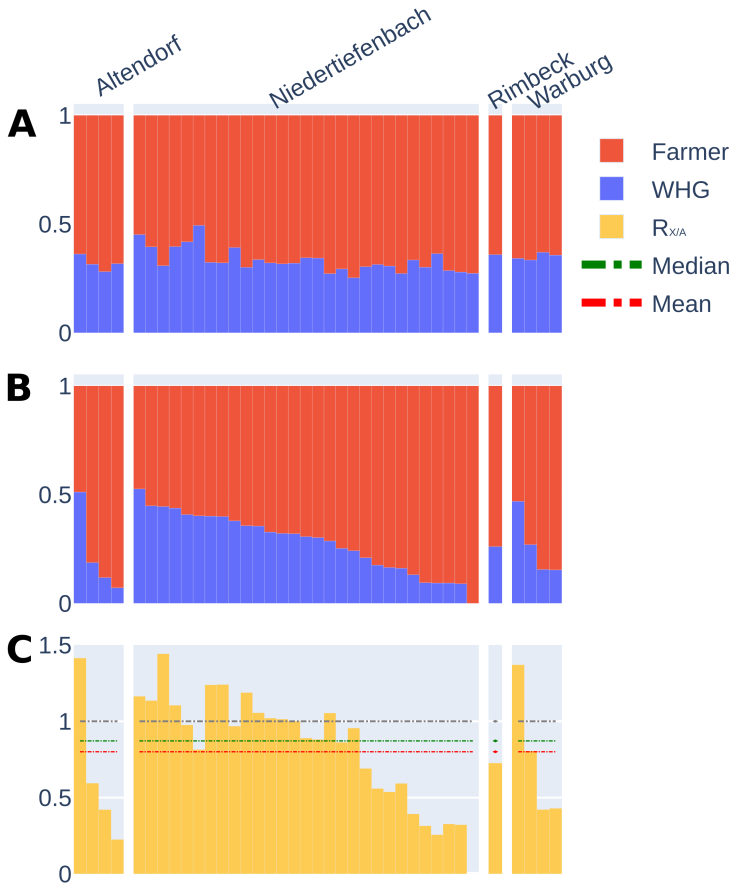


**Figure S8. Results from the sex-bias admixture analyses.** Supervised clustering with two sources: **(A)** using the mean proportions from 1000 resamples with 5388 autosomal SNPs, and **(B)** using 5388 SNPs on the X-chromosome. **(C)** The ratio of X-chromosome to autosomal WHG ancestry ($R_{X/A}$). The grey dashed line refers to the expected value of 1 for an equal contribution of male and female ancestors. The green and red dashed lines refer to the ratio of the median and mean across individuals of X-chromosomal WHG ancestry to individuals of autosomal WHG ancestry.

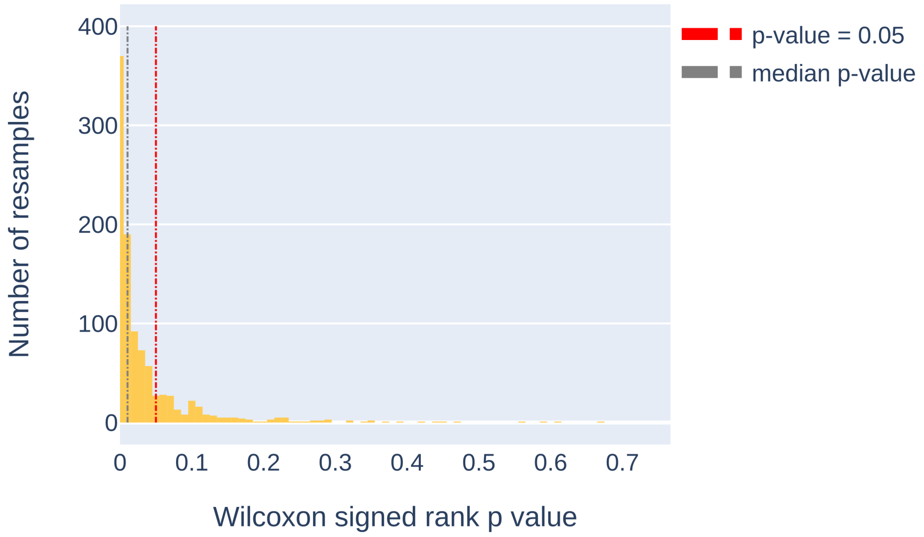


**Figure S9. Statistical test results for sex-biased admixture analysis.** Histogram of p-values for the Wilcoxon sign-rank test performed for the distribution of WHG ancestry on the X-chromosome compared to 1000 autosomal ancestry distributions in late farmers (LF).

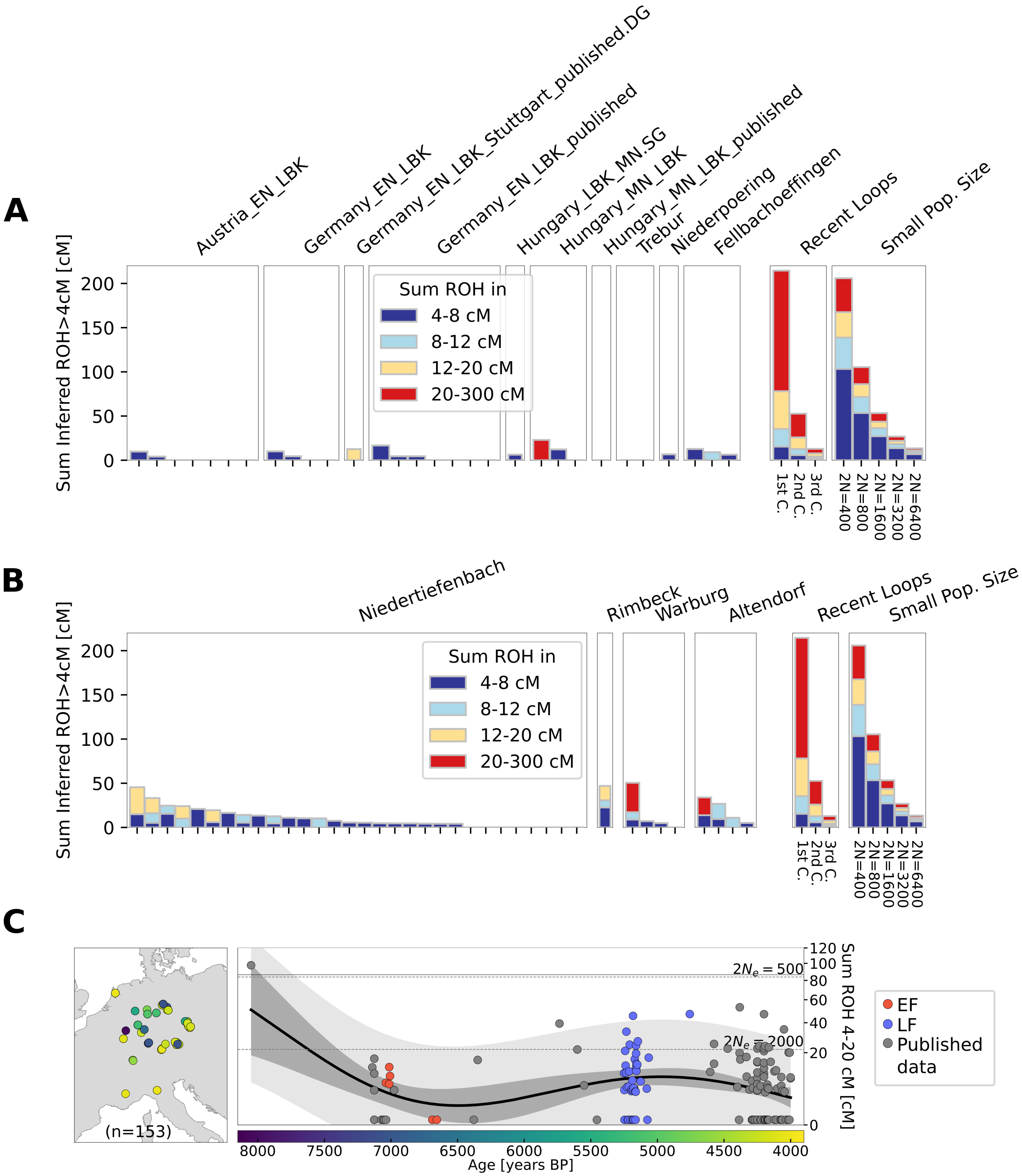


**Figure S10. Estimation of runs of homozygosity (ROH).** Stacked barplots represent total length of ROH, stratified by different length categories in early farmers (EF) and published LBK sites in Central Europe **(A)** and late farmers (LF) **(B)**. Sum of individual ROH (4-20cM) plotted along the timeline of the samples (EF in red, LF in blue, and reference published data in grey) **(C)**. The black solid line shows the average estimates, while the grey areas depict the 95% empirical confidence intervals for both individuals (light grey) and estimated means (dark grey). Only samples with more than 400K covered SNPs were used in the ROH analysis.

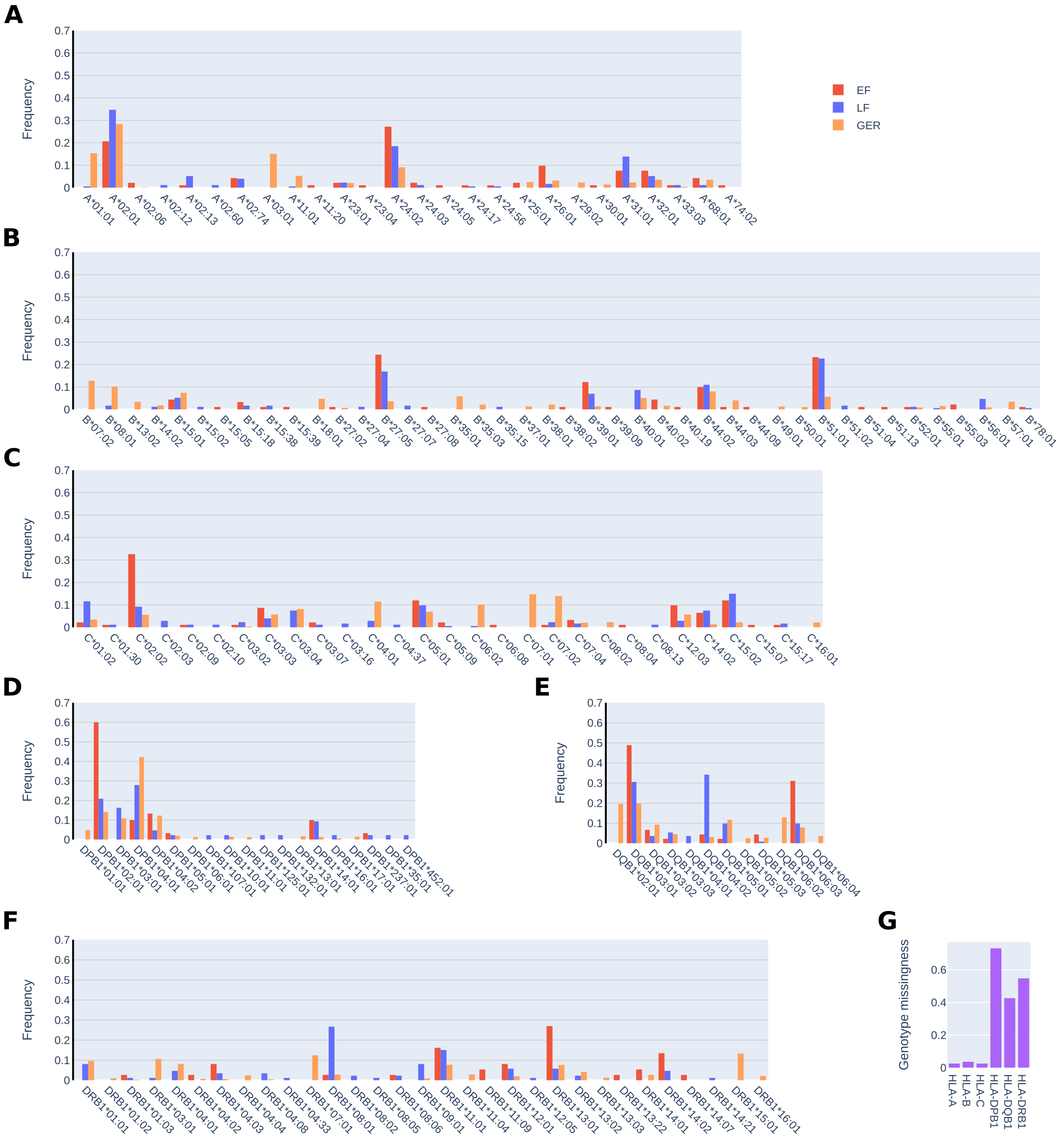


**Figure S11. HLA allele frequencies.** Frequency distribution of HLA-A **(A)**, B **(B)**, C **(C)**, DPB1 (**D**) DQB1 **(E)**, and DRB1 **(F)** alleles in early farmers (EF, blue), late farmers (LF, red) and modern Germans (GER, orange). Only alleles with a frequency above 1% in at least one of the groups is shown in panels (a), (b), (c), (d), (e) and (f). The number of genotyped alleles varied for each locus **(G)**.

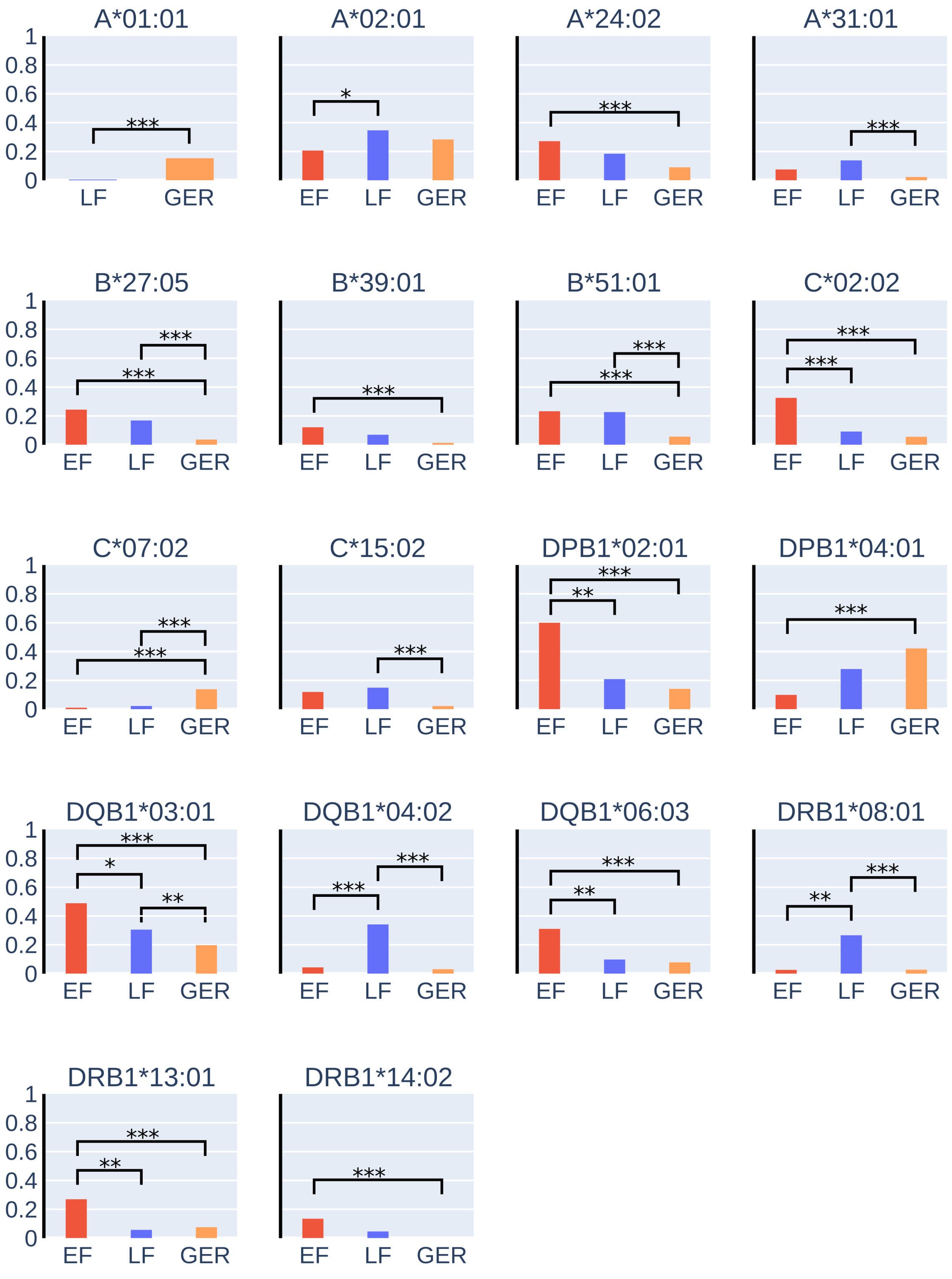


**Figure S12. HLA alleles with significant changes in frequency between early farmers (EF), late farmers (LF) and modern Germans (GER).** Fisher's exact test with multiple test correction was used to estimated p-values (*: $p \leq 0.05$; **: $p \leq 0.01$; ***: $p \leq 0.001$).

**Table S1. List of identified related pairs.**

| Sample pair | Site | # shared SNPs | Mismatches | Mismatch rate | r | r [95% CI lower bound] | r [95% CI upper bound] | Relatedness annotation |
|---|---|---|---|---|---|---|---|---|
| TR124-TR85 | Trebur/Hinkelstein | 3523 | 654 | 0.186 | 0.568 | 0.470 | 0.667 | 1st-degree |
| TR96-TR50 | Trebur/Hinkelstein | 149687 | 29668 | 0.198 | 0.471 | 0.456 | 0.486 | 1st-degree |
| TR96-TR132 | Trebur/Hinkelstein | 31560 | 5971 | 0.189 | 0.541 | 0.508 | 0.574 | 1st-degree |
| TR116-TR76 | Trebur/Hinkelstein | 85276 | 19531 | 0.229 | 0.234 | 0.212 | 0.255 | 2nd-degree |
| TR76-TR83 | Trebur/Hinkelstein | 5685 | 1114 | 0.196 | 0.489 | 0.416 | 0.567 | 1st-degree |
| TR50-TR132 | Trebur/Hinkelstein | 10280 | 2024 | 0.197 | 0.481 | 0.422 | 0.536 | 1st-degree |
| FO130-FO131 | Fellbach Öffingen | 56662 | 11735 | 0.207 | 0.402 | 0.377 | 0.428 | 1st-degree |
| AD165-AD162 | Altendorf | 38855 | 7440 | 0.192 | 0.523 | 0.491 | 0.553 | 1st-degree |
| AD198-AD197 | Altendorf | 14875 | 2994 | 0.201 | 0.447 | 0.398 | 0.496 | 1st-degree |

**Table S2. Comparisons of Fst on the X chromosome and autosomes to explore sex-biased admixture in late farmers (LF).**

| Comparison | Fst (autosomes) | Fst (chrX) | Q |
|---|---|---|---|
| LF-WHG | 0.042 | 0.064 | 0.634 |
| LF-Anatolia | 0.025 | 0.033 | 0.755 |

**Table S3. HLA alleles exhibiting statistically significant and substantial shifts ($p \leq 0.05$, absolute frequency difference ≥ 10%) between early farmers (EF) and late farmers (LF) or either Neolithic groups and modern Germans (GER).** The p-values were calculated with the Fisher's exact test and corrected for multiple testing using two stage Benjamini and Hochberg step-up FDR-controlling procedure (TSBH-FDR).

| HLA allele | Test | Freq. difference | p-value |
|---|---|---|---|
| A*02:01 | EF vs LF | 0.14 | 2.47E-02 |
| C*02:02 | EF vs LF | 0.23 | 1.50E-05 |
| DPB1*02:01 | EF vs LF | 0.39 | 1.64E-03 |
| DQB1*03:01 | EF vs LF | 0.18 | 4.24E-02 |
| DQB1*04:02 | EF vs LF | 0.3 | 6.42E-05 |
| DQB1*06:03 | EF vs LF | 0.21 | 3.67E-03 |
| DRB1*08:01 | EF vs LF | 0.24 | 1.60E-03 |
| DRB1*13:01 | EF vs LF | 0.21 | 2.62E-03 |
| A*24:02 | EF vs GER | 0.18 | 1.59E-06 |
| B*27:05 | EF vs GER | 0.21 | 9.25E-12 |
| B*39:01 | EF vs GER | 0.11 | 1.80E-07 |
| B*51:01 | EF vs GER | 0.18 | 9.68E-08 |
| C*02:02 | EF vs GER | 0.27 | 2.55E-14 |
| C*07:02 | EF vs GER | 0.13 | 4.92E-05 |
| DPB1*02:01 | EF vs GER | 0.46 | 3.19E-08 |
| DPB1*04:01 | EF vs GER | 0.32 | 3.88E-04 |
| DQB1*03:01 | EF vs GER | 0.29 | 3.08E-05 |
| DQB1*06:03 | EF vs GER | 0.23 | 1.65E-05 |
| DRB1*13:01 | EF vs GER | 0.19 | 4.97E-04 |
| DRB1*14:02 | EF vs GER | 0.14 | 6.68E-14 |
| A*01:01 | LF vs GER | 0.15 | 1.29E-10 |
| A*31:01 | LF vs GER | 0.12 | 3.11E-11 |
| B*27:05 | LF vs GER | 0.13 | 5.58E-11 |
| B*51:01 | LF vs GER | 0.17 | 1.03E-12 |
| C*07:02 | LF vs GER | 0.12 | 8.84E-07 |
| C*15:02 | LF vs GER | 0.13 | 4.89E-13 |
| DQB1*03:01 | LF vs GER | 0.11 | 7.12E-03 |
| DQB1*04:02 | LF vs GER | 0.31 | 2.09E-27 |
| DRB1*08:01 | LF vs GER | 0.24 | 4.68E-15 |

**Table S4. Most common HLA alleles (frequency ≥ 10%) in modern Germans (GER) compared to their frequencies in Neolithic groups (early farmers (EF) and late farmers (LF)).**

| HLA allele | Frequency in EF | Frequency in LF | Frequency in GER |
|---|---|---|---|
| A*01:01 | 0 | 0.006 | 0.154 |
| A*02:01 | 0.207 | 0.347 | 0.284 |
| A*03:01 | 0 | 0 | 0.151 |
| B*07:02 | 0 | 0 | 0.128 |
| B*08:01 | 0 | 0.017 | 0.102 |
| C*04:01 | 0 | 0.029 | 0.115 |
| C*06:02 | 0 | 0.006 | 0.101 |
| C*07:01 | 0 | 0 | 0.147 |
| C*07:02 | 0.011 | 0.023 | 0.139 |
| DPB1*02:01 | 0.6 | 0.209 | 0.142 |
| DPB1*03:01 | 0 | 0.163 | 0.108 |
| DPB1*04:01 | 0.1 | 0.279 | 0.422 |
| DPB1*04:02 | 0.133 | 0.047 | 0.123 |
| DQB1*02:01 | 0 | 0 | 0.196 |
| DQB1*03:01 | 0.489 | 0.306 | 0.198 |
| DQB1*05:01 | 0.022 | 0.099 | 0.117 |
| DQB1*06:02 | 0 | 0 | 0.129 |
| DRB1*03:01 | 0 | 0.012 | 0.107 |
| DRB1*07:01 | 0 | 0 | 0.125 |
| DRB1*15:01 | 0 | 0 | 0.133 |
